# Differentiating the Roles of Metabolic Similarity and Ionic Coupling in Determining the Beta Hub Cell Phenotype

**DOI:** 10.1101/2025.10.27.684924

**Authors:** Ingvild S. Devold, Hanne Rokstad, Jazmin Velazquez, Shane Alexander Browne, Andrew G. Edwards, Vira Kravets

**Affiliations:** Department of Numerical Analysis and Scientific Computing, Simula Research Laboratory, NO; Department of Physics, Norwegian University of Science and Technology, NO; Department of Bioengineering, University of California Berkeley, USA; Department of Mechanical and Aerospace Engineering, University of California San Diego, USA; Department of Computational Physiology, Simula Research Laboratory, NO; Department of Bioengineering & Department of Pediatrics, University of California San Diego, USA

## Abstract

Pancreatic beta cells regulate circulating glucose levels by releasing insulin. Beta cells transduce elevated blood glucose into electrical activity through an intrinsic cascade of metabolic and electrophysiologic responses. These responses are synchronized across the electrically connected network of beta cells, such that insulin release is also relatively synchronous. Despite their coordinated behavior, individual beta cells exhibit significant functional heterogeneity. This heterogeneity is thought to provide cells with specific functional phenotypes the ability to control the activity of the broader network. Hub cells, identified by their synchronized [Ca^2+^] activity, are one such functional subpopulation, and are believed to orchestrate the second, oscillatory phase of insulin release. However, it remains unclear whether the cell-autonomous characteristics of hub cells, such as their metabolic activity and electrophysiologic properties, are more important than their network characteristics (i.e. gap junctional coupling) for their ability to influence broader network activity. In this study, we investigate the roles of intrinsic metabolic and electrophysiologic properties and ionic coupling in determining the beta hub cell phenotype. Using a computational islet model of 1,000 beta cells, our analysis revealed that both intrinsic metabolic properties and structural coupling via gap junctions are crucial for determining the hub cell phenotype. After investigating the intrinsic coupling conductance of neighboring cells and the number of structural (direct electrical) links as independent contributors to the hub cell’s local electrical coupling, we find that the number of cells to which a beta cell is directly coupled may be a key determinant of its propensity to serve as a hub cell in the model. As predicted for this subpopulation, we also demonstrate that decoupling hub cells impairs the functional connectivity of the entire network. Our findings indicate the importance of both autonomous cellular dynamics and non-autonomous structural coupling for the hub cell phenotype. These insights help build a fundamental understanding of hub cells, which in turn may contribute to identifying potential approaches to preserve or improve beta cell function and thereby manage the progression of diabetes.

## 4.1 Introduction

Beta cells in the pancreatic islets regulate blood glucose levels through insulin secretion. These cells respond to changes in glucose concentrations by generating electrical signals that trigger insulin release, and their dysfunction is linked to diabetes. Although the responses of beta cells often appear highly synchronized, these cells are functionally heterogeneous. Different subpopulations have been proposed to play distinct roles in modulating islet function. “First responder” beta cells have been hypothesized to drive the first phase of insulin release by reacting most quickly to glucose elevation and thereby initiating the electrical activity that in turn drives increased [Ca^2+^] and insulin secretion[1, 2]. Experimental and computational studies have also identified “hub” cells, which are distinguished by their highly synchronized [Ca^2+^] activity during the second oscillatory phase of insulin release and thought to be largely responsible for coordinating second phase activity of the islet[3, 4, 5].

In response to glucose, increased glycolytic flux and ATP production cause ATP-sensitive potassium channels (K_ATP_) to close, leading to membrane depolarization. This triggers the opening of voltage-gated calcium channels, resulting in elevated intracellular [Ca^2+^], which in turn stimulates insulin release. Importantly, each beta cell is not an isolated functional entity, but one functional component of a large network of beta cells that is electrically coupled through connexin 36 (Cx36) gap junctions [6]. This gap junctional coupling allows the cells to synchronize their electrical activity and generate a pulsatile insulin release, coordinated across the islet. In combination, this signaling cascade and structural connectivity causes beta cell and islet function to be influenced by both the intrinsic, autonomous properties of each cell and non-autonomous properties arising from the cell’s role in a network connected by gap junctions.

These complex interactions are challenging to study experimentally, but they lend themselves to computational modeling. To better understand the relative importance of autonomous and non-autonomous factors, we apply a computational model of a mouse pancreatic beta cell network [7, 8, 2, 5]. This model, an extension of the single beta cell Cha-Noma model [9], simulates a network of 1000 beta cells, electrically coupled through gap junctions [7]. The model allows us to simulate calcium dynamics, and thereby identify hub cells via correlation analysis, analogously to experimental approaches [10, 3]. A key advantage of this computational approach is the opportunity to systematically investigate the role of intrinsic properties of beta cells, such as metabolic parameters, or their network properties, such as gap junction coupling, for determining overall islet function.

In this study, we aim to understand how both intrinsic cell dynamics and communication through gap junctions determine hub cell properties within a beta cell network. By simulating islets with variable parameters and decoupling different subpopulations on the basis of those parameters or of their function within the network, we can explore how intrinsic cell dynamics and gap junctional coupling influence the overall function of pancreatic islets. Proper understanding of these interactions is essential in preserving and improving beta cell function to potentially develop targeted therapies preventing the progression of diabetes.

### 4.2 Methods

#### Coupled Cha-Noma beta cell model

We used a previously introduced computational model of a heterogeneous network of beta cells [7], implemented in C++. The model builds upon the Cha-Noma model [9, 11], which describes the electrophysiological properties of a single mouse beta cell, including ion channels, transporters, and [Ca^2+^] dynamics (see Figure 4.1). This is extended to a network of *N* = 1000 cells, with between 1 and 12 gap junction connections per cell [5]. The system is described by a set of ordinary differential equations (ODEs), where the membrane potential *V*_*i*_ of each cell *i* is governed by

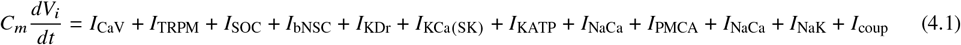

and solved for in each time step. Here, the coupling current,

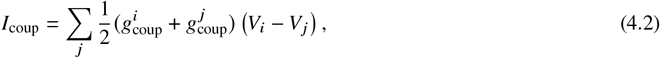

models the current through the cell’s Cx36 gap junctions and depends on the voltage difference between the connected cell pairs (*i, j*) and both their intrinsic coupling conductances, *g*_coup_.

**Fig. 4.1.**
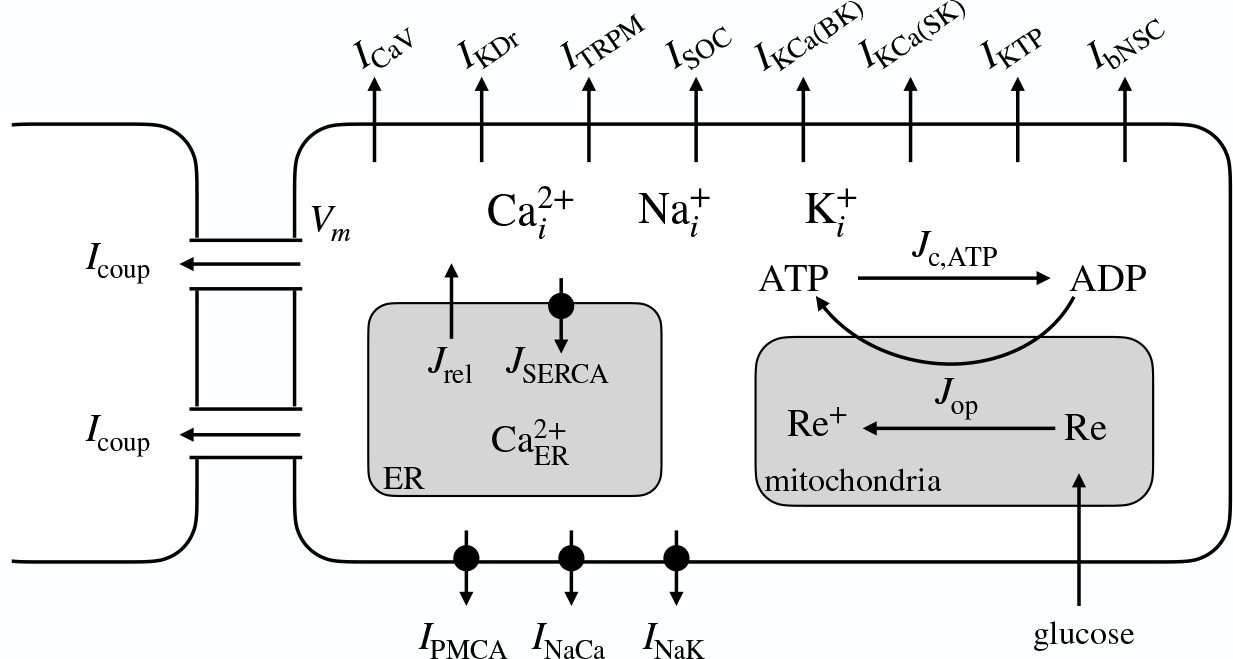
Schematic diagram of the coupled Cha-Noma model. Figure recreated with modification from [9].

#### 4.2.1 Cellular heterogeneity from randomized parameter values

Heterogeneity was introduced to the model by assigning randomized parameter values to cells based on previously established distributions from experiments [7]. Table 4.1 shows the distributions used for three of the parameters. The parameter assignments were seeded, and for our experiments, we ran repeated simulations for different seeds, in an attempt to mimic the variability found in experimental islets and to enhance the statistical robustness of the analysis.

**Table 4.1.** A selection of probability distributions used when assigning randomized parameter values to cells in the computational beta cell network model [5].

| Parameter | Symbol | Distribution | Mean | SD | Unit |
| --- | --- | --- | --- | --- | --- |
| Glycolysis rate | $k_{\text{glyc}}$ | Normal | 0.126 | 0.0315 | $\text{ms}^{-1}$ |
| Coupling conductance | $g_{\text{coup}}$ | Gamma(4,4) | 0.12 | 0.06 | nS |
| $K_{\text{ATP}}$ channel conductance | $g_{\text{KATP}}$ | Normal | 2.31 | 0.57 | pA/mV |

#### 4.2.2 Driving glucose curve

The primary driver of the simulation was a prescribed glucose curve [G] (*t*). We used a simple step function for this purpose, starting with an initial glucose concentration of [G] = 2.0 mM for the first 20 seconds. After this period, the glucose concentration was increased to a chosen maximum level (e.g. [G] _max_ = 12.0 mM), which was maintained for the remainder of the simulation. We tested [G] _max_ ∈ {2, 4, 6, 8, 10, 12} mM, and ran analysis on the [G] _max_ = 12 mM case since this consistently produced [Ca^2+^] oscillations across seeds.

#### 4.2.3 Defining the functional network from Ca^2+^ activity

Among the simulation outputs were the [Ca^2+^] time series data for each of the *N* = 1000 beta cells. These time series were used to construct the islet functional network by identifying highly synchronized cells, as described in [5]. First, we extracted a time window containing the second, oscillatory phase. For simulations with duration 900 s, we used the period from 100 s to the end of the simulation. With the [Ca^2+^] data from this time window, we computed the correlation matrix using MATLAB’s corr() function, which provides the Pearson correlation coefficients,

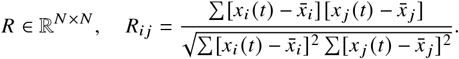

Next, we applied a threshold *R*_th_ to identify synchronized cell pairs. Cell pairs with correlation coefficients above this threshold were considered functionally linked. The resulting adjacency matrix *A*, with entries

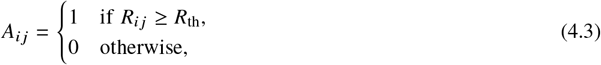

defined the beta cell functional network. We chose the threshold *R*_th_ = 0.99975 to yield a power law distribution of network degree, as suggested by experimental studies [12, 3].

#### 4.2.4 Analysis

##### 4.2.4.1 Identifying hub cells from network degree

The degree of a node in a network refers to its number of edges. In the context of the beta cell functional network, this corresponds to the number of cells a given cell is highly synchronized with, and therefore likely to be functionally coupled with. In this simulation framework, the only direct means of coupling is gap junctional. We identified hub cells as those whose degree exceeded 65% of the maximal degree observed for any cell in the network (deg_*th*_ = 0.65). This criterion resulted in a proportion of hub cells consistent with the previously suggested range of 1-10% [3].

##### 4.2.4.2 Comparative analysis of hubs and followers

With the population divided into hub and follower (non-hub) cells, means and standard errors (SEM) were used to visually compare parameter values, unless otherwise specified. Statistical inference was performed via two-sample t-test, using MATLAB’s ttest2() function for comparison of hubs vs. followers. All t-tests were performed at the seed level, such that the mean parameter values for hubs and followers from each seed constituted the independent observations subjected to t-tests. In addition to comparing the assigned randomized parameters, we calculated the *total* gap junction coupling conductance as suggested in [5],

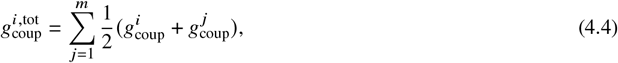

summing the average coupling conductances of the cell and its physically linked neighbors (*j* = 1, …, *m*). This parameter incorporates both the number of gap junctions of a given cell, its intrinsic coupling conductance, and the intrinsic coupling conductances of its connected cells.

###### Decoupling beta cell subpopulations

Following the original islet simulations and subsequent analysis, we re-ran the simulations after decoupling specific subsets of beta cells. For example, all cells in the top 10% of the coupling distribution. In this process we simulated removal of those cells’ gap junctions by excluding their coupling current contributions (Equation 4.2) from the ODE system. In these decoupled simulations, the decoupled subpopulation was also excluded from the subsequent analysis. To ensure a fair comparison between simulated decoupling experiments, we consistently decoupled a fixed number of cells. This approach prevented variations in the size of the decoupled subgroups. Finally, the decoupling of a random group of cells served as a control, allowing us to isolate the effects of decoupling specific subpopulations from the more general effect of the decoupling itself.

### 4.3 Results

To analyze the relative importance of functional similarity and structural network connectivity to the beta hub cell phenotype, we analyzed a simulated beta cell network using an extension of the Cha-Noma model [9]. The model incorporates both the main ion channels affecting the beta cell membrane potential and the electrical coupling between cells through Cx36 gap junction channels. Cellular heterogeneity was implemented through randomized parameters (Figure 4.2A), assigned according to probability distributions based on experimental data [7]. The islet was stimulated by a [G] _max_ = 12.0 mM maximum glucose concentration from 20 s to 900 s. We repeated the simulation for 8 different islets, each with different randomized parameter values. This produced fast (¡ 2 min) and highly synchronized Ca^2+^ oscillations for all simulated islets. Figure 4.2B shows a representative calcium response of one simulated islet, and Figure 4.2C highlights the tight coupling of the network at two time points – at a peak and a nadir of the calcium response.

**Fig. 4 2.**
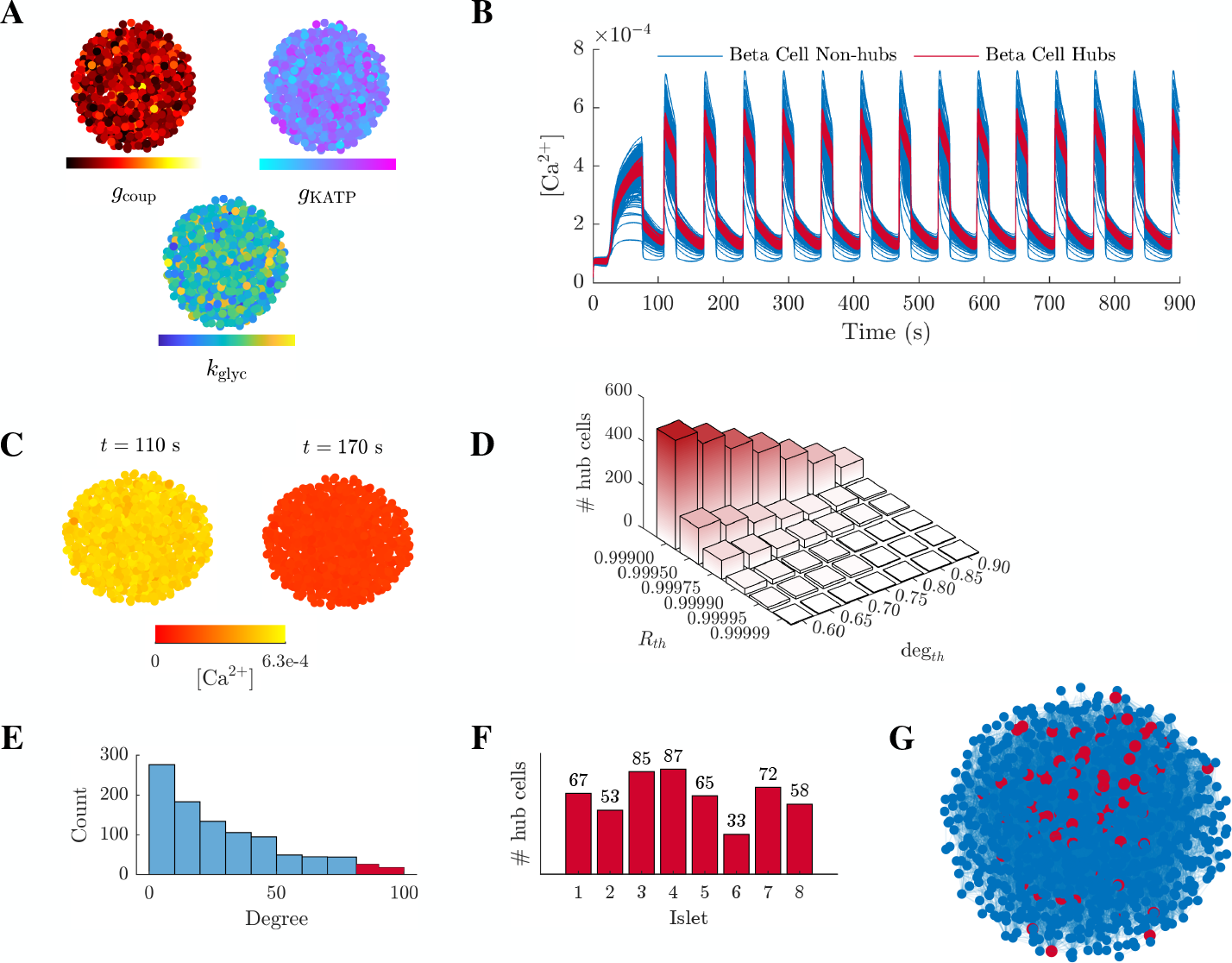
From structural to functional beta cell network. (**A**) The 1000-cell beta cell network false-colored by parameter values (gap junction coupling conductance *g*_coup_, *K*_ATP_ channel conductance *g*_KATP_, and glycolysis rate *k*_glyc_). (**B**) Calcium response from the islet shown in (A), simulated at [*G*] _max_ = 12 mM for 900 s, with hub cells highlighted in red. (**C**) False-color map displaying the beta cell network calcium concentration at two time points, near a peak (*t* = 110 s) and a nadir (*t* = 170 s). (**D**) Number of identified hub cells for different correlation (*R*_*th*_) and network degree (deg_*th*_) thresholds. (**E**) Degree distribution of the functional network. Hub cells with normalized degree exceeding 0.65 highlighted in red. (**F**) Number of identified hubs for eight different simulated islets, using the *R*_*th*_ = 0.99975 and deg_*th*_ = 0.65 criteria. (**G**) Beta cell functional network with identified hubs highlighted in red.

### 4.3.1 Two thresholds determine the number of identified hub cells

We examined the effect of two thresholds on the number of identified hub cells (Figure 4.2D). Our analysis confirmed that both the correlation threshold *R*_*th*_ and the degree threshold deg_*th*_ impact the number of hub cells identified, with larger thresholds resulting in fewer hubs. Moreover, the correlation threshold *R*_*th*_ directly affects the degree distribution of the functional network. Figure 4.2E displays the power-law-like distribution obtained using a value *R*_*th*_ = 0.99975. Using the *R*_*th*_ = 0.99975 and deg_*th*_ = 0.65 criteria, we obtained hub cell proportions between 3% and 9% for the different simulated islets (Figure 4.2F). This is within the 1%-10% interval previously advocated for based on experiments [3]. Finally, Figure 4.2G shows the spatial distribution of the identified hub cells in the functional network for one simulated islet. We found no discernible pattern in the spatial organization of the hub cells. Instead, they appeared to be randomly scattered across the network.

### Simulations of Ca^2+^ oscillations reveal that both metabolic activity and structural coupling are key in differentiating beta cell hubs

Next, we analyzed 8 simulated islets with different randomized parameter sets to investigate systematic differences between hub and follower beta cell subpopulations. The glucokinase activity, quantified by the glycolysis rate *k*_glyc_, serves as an indicator of the cell’s metabolic activity. We found that *k*_glyc_ was significantly higher in hub cells than follower cells (mean ± SD 0.14 ± 0.0022 ms^−1^ vs. 0.12 ± 0.0009 ms^−1^, *p* < 0.001) (Figure 4.3A). The primary point of coupling between metabolic and electrical activity in beta cells is the ATP-sensitive potassium channel. The conductance of these channels (*g*_KATP_), which corresponds to the number of *K*_ATP_ channels, is expected to impact cell dynamics. We observed that *g*_KATP_ was slightly lower in hub cells compared to follower cells (mean ± SD 2.27 ± 0.05 pA/mV vs. 2.31 ± 0.02 pA/mV, *p* = 0.03) (Figure 4.3B). These results suggest that, within this model, metabolic activity plays a more important role than the number of *K*_ATP_ channels in determining the hub cell phenotype. This finding aligns with previous research [5].

**Fig. 4.3.**
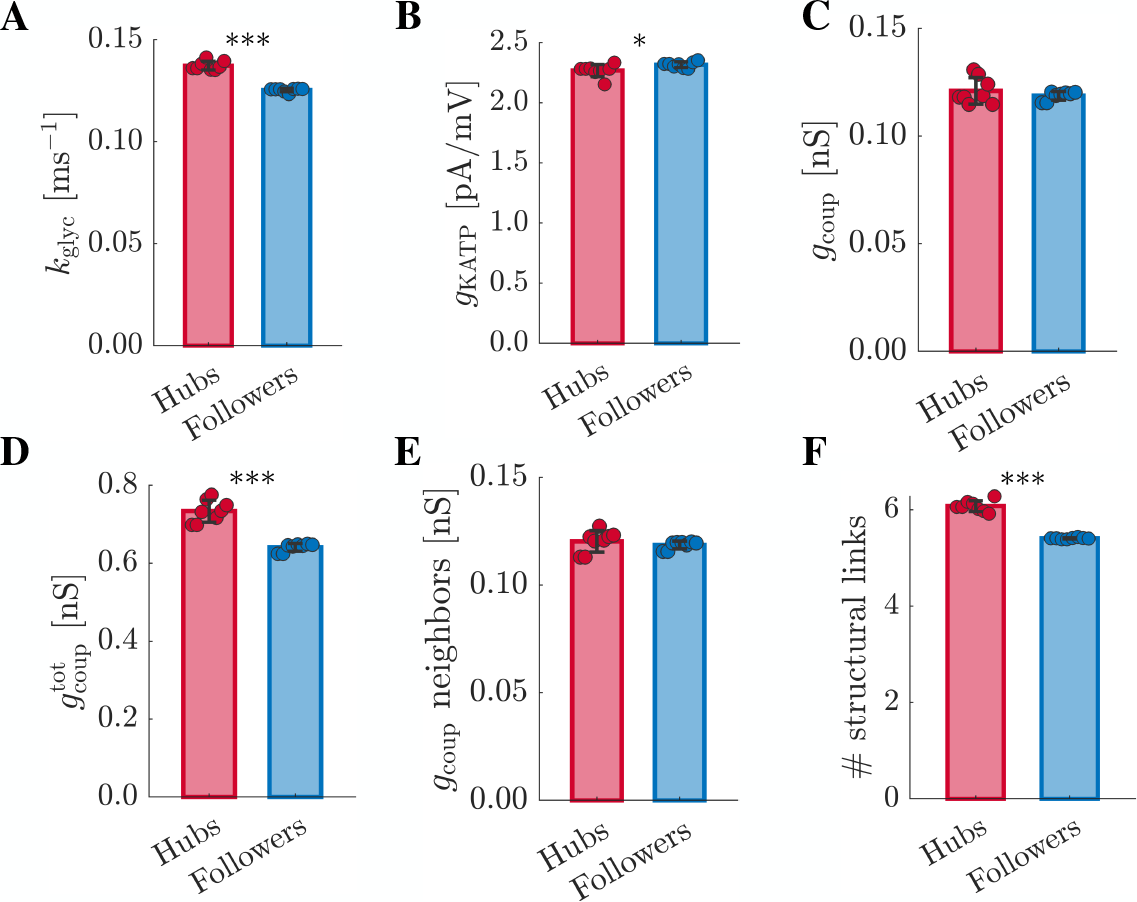
Analysis of parameter differences between the hub and follower beta cell subpopulations. Significance was determined by paired t-tests. ^*^*p* < 0.05, ^**^*p* < 0.01, ^***^*p* < 0.001. Data points represent average values of each simulated islet, and error bars represent SD. (**A**) Mean parameter value of glycolysis rate (*k*_glyc_) averaged over the hub and follower (non-hub) cell subpopulations (*n* = 8 seeds). (**B**) As in (A) for *K*_ATP_ conductance (*g*_KATP_). (**C**) As in (A) for coupling conductance (*g*_coup_). (**D**) As in (A) for total coupling conductance 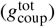. (**E**) As in (A) for mean coupling conductance of structurally linked cells. (**F**) As in (A) for number of structural links.

Intuitively, it is reasonable to expect that structural coupling affects network synchrony and therefore high structural coupling may be a characteristic feature of beta hub cells. Still, we observed that intrinsic coupling conductance (the *g*_coup_ attributable only to each cell, rather than the cell and its direct connections) did not differ significantly between hub and follower cells (mean ± SD 0.121 ± 0.006 nS vs. 0.118 ± 0.002 nS, *p* = 0.33) (Figure 4.3C). However, comparing the total gap junction coupling conductance (Equation 4.4), we observed that this was significantly higher for hub cells than followers (mean ± SD 0.73 ± 0.03 nS vs. 0.64 ± 0.01 nS, *p* < 0.001) (Figure 4.3D). This suggests that both the intrinsic coupling conductance of the cell, and the coupling conductances of cells in its direct network (those coupled to it) may act in concert to make the cell highly synchronized. We next decomposed how those coupled neighbors influence the hub cell phenotype. We found that the mean intrinsic coupling conductance of a hub cell’s structurally linked neighbors was not different (mean ± SD 0.120 ± 0.005 nS vs. 0.119 ± 0.002 nS, *p* = 0.36) (Figure 4.3E) from that of the follower cell’s. Instead, the number of structural links was higher for hub cells than for followers (mean ± SD 6.1 ± 0.11 vs. 5.4 ± 0.02, *p* < 0.001) (Figure 4.3F). Together, these observations indicate that the number of cells to which a beta cell is directly coupled may be a key determinant of its propensity to serve as a hub cell in the model.

**Fig. 4. 4.**
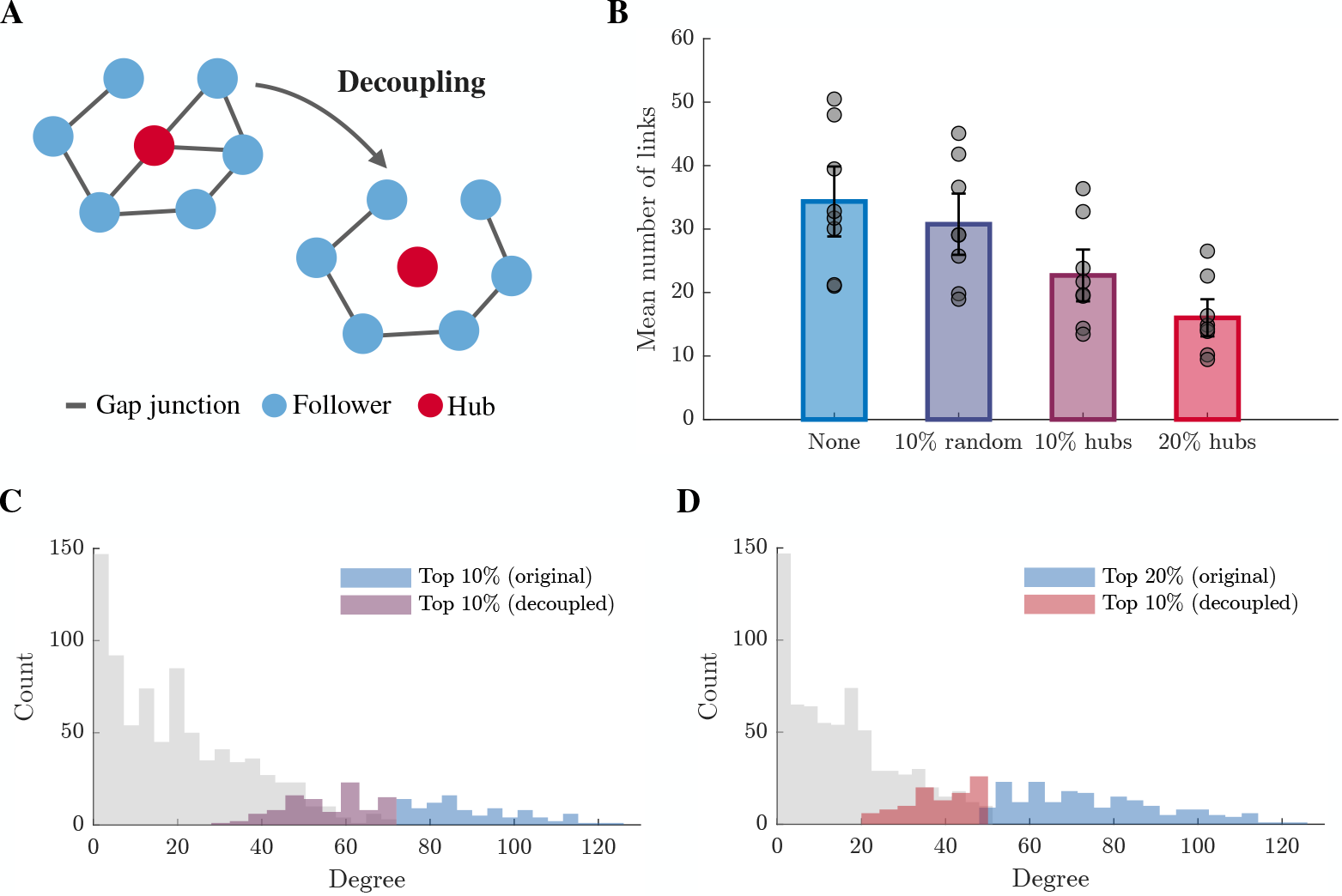
Analysis of decoupled islet simulations. (**A**) Schematic of the decoupling process, where the gap junctions of hub cells are removed. (**B**) Mean number of functional links for different decoupled subpopulations. From left to right, the following groups were decoupled: None; 100 random cells; the 100 most functionally linked cells; the 200 most functionally linked cells. The bars indicate the mean degree across all simulated islets (*n* = 8 seeds), whereas the data points represent individual islets. Error bars indicate SEM. (**C**) Degree distribution of the original simulated islet, with cells selected for decoupling (“Top 10% (original)”) and the new 100 most functionally linked in the decoupled islet (“Top 10% (decoupled)”) indicated. (**D**) As in (C), but decoupling the 200 most functionally linked cells.

### Decoupling beta cell subpopulations reveals impaired islet connectivity and potential compensatory mechanisms

To investigate the importance of selected beta cell subpopulations to the total islet synchronization, we ran decoupled simulations on eight different simulated islets. This entailed removing the gap junctions of selected subsets of cells from the simulation and excluding those cells from the subsequent analysis (Figure 4.4A). We compared the effect of decoupling three different groups of cells on the network connectivity, measured by the mean number of functional links (Figure 4.4B). First, we decoupled 100 random cells, in an attempt to study the isolated effect of the decoupling itself. This slightly reduced the mean number of functional links, from 34.3 ± 5.5 to 30.8 ± 4.8 (mean ± SEM) (Figure 4.4B). Decoupling the 10% most functionally linked cells substantially reduced the average connectivity to 22.7 ± 4.1. Finally, decoupling the 20% most functionally linked had the strongest effect, resulting in an average number of functional links of 16.0 ± 2.9 in the remaining coupled islet.

With decoupling, we aimed to simulate the effect of cell ablation. Although the total islet synchronization was impaired after decoupling, the islet remained tightly coupled, raising the question of whether other cells can take on the roles of the pacemaker hubs. To answer this, we identified the 100 most functionally linked cells in the decoupled simulations and examined where those cells belonged in the degree distribution of the original islet. After decoupling the 10% most linked cells, the new 10% most linked group of cells appeared to be on the upper end of the degree spectrum of the remaining cells (Figure 4.4C). Similarly, after decoupling the top 20%, the replacement hubs appeared to be in the next degree percentiles of the original islet (Figure 4.4D). Thus, it is likely that the hierarchy of functional coupling is well described by the degree distribution, and that any cell’s position within that distribution (hierarchy) is not markedly altered by removing a reasonably small fraction of highly coupled cells.

## 4.4 Discussion

In this study, we integrated computational modeling with principles of biological heterogeneity to investigate the defining characteristics of hub cells in pancreatic beta cell networks. Our findings indicate that the hub cell phenotype arises from a combination of metabolic properties and structural connectivity. This dual dependency emphasizes the importance of both autonomous cellular functions and direct cell-cell electrical coupling in regulating insulin secretion and maintaining glucose homeostasis, contributing to novel insights into islet physiology.

We demonstrate that hub cells exhibit significantly higher glycolytic rates (*k*_glyc_) compared to follower cells, suggesting that intrinsic metabolic activity is a critical determinant of hub cell functionality. This aligns with previous studies that identified metabolic heterogeneity as a driver of functional differences within beta cell populations [4]. The higher glycolytic activity in hub cells likely facilitates their ability to coordinate the second phase of insulin release, a process essential for maintaining normoglycemia. This suggests that targeting metabolic pathways specific to hub cells could offer a novel approach to enhancing islet function in diabetic conditions.

The role of structural connectivity, via gap junctions, is also essential for defining the hub cell phenotype. Our simulations indicate that hub cells have significantly higher total coupling conductance 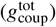, enhancing synchronized activity across the islet. We also decompose this effect by investigating the intrinsic coupling conductance of neighboring cells and the number of structural (direct electrical) links as independent contributors to the hub cell’s local electrical coupling. These analyses demonstrate that the difference in 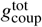 likely stems from the number of cells to which they are directly coupled rather than the degree or coupling conductance of those connected cells. The decoupling experiments emphasize that disrupting gap junctions, specifically in the highly connected hub cells, leads to a decline in coordination across the islet. This highlights the importance of hub cells in ensuring effective propagation of electrical signals and thereby coordination of insulin release.

While the computational model offers valuable insights, it also has limitations. A challenge with the current model is its extremely tight coupling. The [Ca^2+^] time series are all highly correlated, illustrated by the large threshold (*R*_*th*_ = 0.99975) required to achieve a power-law degree distribution. This tight coupling makes it difficult to discern clear effects of any manipulation on synchronization or functional organization of the islet. When all cells are that highly correlated, the intricate dynamics and potential heterogeneity within the islet may be obscured. This underscores the importance of exploring various decoupling strategies to reveal underlying patterns and to better understand the functional connectivity within the islet. By selectively decoupling specific groups of cells, we aim to expose the subtle nuances of cellular interactions that are otherwise concealed in a highly coupled system. This also highlights the sensitivity of the islet analysis to the manually chosen threshold. It is essential to solidify the choice of this threshold, and additional experiments should be considered for validation.

Another challenge arises from the significant variability between simulated islets. When different seeds are used to assign randomized parameters to cells, this variability leads to substantial differences in islet behavior and organization, making it difficult to draw consistent conclusions. This issue was especially apparent in the decoupled simulations, where the mean number of functional links varied considerably between seeds. Each set of random parameters can create unique patterns of synchronization and connectivity, complicating generalization. To address this, robust statistical methods and multiple simulations are necessary to capture the range of possible outcomes and to reflect the variability found in real-world islets. Analyzing a wide array of simulations helps us understand the variability and identify consistent patterns.

In the future, extending the model to include pathological conditions, such as diabetes, could provide valuable insights into how hub cell functionality changes with disease states. Furthermore, our results suggest that hub cells could serve as biomarkers for islet health, with potential applications in the early detection of beta cell failure. The integration of computational and experimental approaches will be essential in further explaining the role of hub cells and their potential as targets for preserving islet function.

## 4.5 Conclusion

Our study highlights the critical role of both autonomous and non-autonomous parameters in defining the beta hub cell phenotype. Employing the extended Cha-Noma model, we established that autonomous metabolic factors, such as glycolysis rate, are essential in phenotype differentiation. Non-autonomous factors, including total gap junction coupling conductance and the number of structural links, also play a significant role in hub cell identification. Hub cells act as critical coordinators within the beta cell network, with their functionality emerging from a delicate balance between metabolic activity and structural connectivity. The seemingly random spatial distribution of hub cells suggests that location is not a significant factor, at least in our model of a beta cell islet. Lastly, our decoupling experiments emphasize the importance of hub cells in maintaining network synchronization. Decoupling these highly coupled cells leads to significant disruptions. Still, the islet remains highly synchronized, suggesting that other cells may partially compensate for the missing hub cells, thereby ensuring continued functionality despite disruptions. These findings contribute to the understanding of islet physiology and suggest new avenues for diagnostics and potential therapeutic interventions in diabetes.

